# The Central Nervious System Acts as a Transducer of Stress-Induced Masculinization through Corticotropin-Releasing Hormone B

**DOI:** 10.1101/415554

**Authors:** DC Castañeda Cortés, LF Arias Padilla, VS Langlois, GM Somoza, JI Fernandino

**Affiliations:** Laboratorio de Biología del Desarrollo - Instituto Tecnológico de Chascomús. INTECH (CONICET-UNSAM), Argentina; Institut national de la recherche scientifique (INRS) - Centre Eau Terre Environnement, Quebec, Canada; Laboratorio de Ictiofisiología y Acuicultura - INTECH (CONICET-UNSAM), Argentina

**Keywords:** Environmental stress, CRH, Masculinization, CRISPR/Cas9, Medaka

## Abstract

Exposure to environmental stressors during early development has important implications for rescheduling many cellular and molecular mechanisms. In some fish species, environmental stressors, like high temperatures (HT), cause an increase in cortisol levels. In turn, this mechanism induces sex reversal of genotypic females, overriding genetic factors related to development of the gonad. However, the involvement of the brain in this process is not well clarified. In the present work, we investigated the mRNA levels of corticotropin-releasing hormone b (*crhb*) and its receptors (*crhr1* and *crhr2*), and found out that they were up-regulated at HT during the critical period of gonadal sex determination in medaka (*Oryzias latipes*), i.e., when the gonadal primordium is sexually labile. In order to clarify their roles in sex reversal, biallelic mutants for *crhr1* and *crhr2* were produced by CRISPR/Cas9 technology. Remarkably, biallelic mutant of both *loci* (*crhr1* and *crhr2*) did not undergo female-to-male sex reversal upon HT exposition, whereas mutants for either *crhr1* or *crhr2* showed partial, or intersex phenotypes, suggesting that both *crh* receptors are required for HT-induced masculinization. Inhibition of this process in double *crhr*s mutants could be successfully rescued through the administration of the downstream effector of the hypothalamic-pituitary interrenal axis, the cortisol. Taken together, these results revealed for the first time the participation of the central nervous system acting as a transducer of masculinization induced by thermal stress.

## INTRODUCTION

As a general trend, the response of the neuroendocrine system to environmental stressors produces the elevation of the hypothalamic corticotropin-releasing hormone (CRH). CRH in turn stimulates the secretion and release of adrenocorticotropic hormone (ACTH) from the pituitary gland (Aguilera & Liu, 2012; Kovacs, 2013), regulating cortisol levels through the adrenal gland (Mommsen, Vijayan, & Moon, 1999). This axis is known as hypothalamic-pituitary-adrenal (HPA) or the -interrenal (HPI) in tetrapods and fish, respectively. In this last group, two *crh* ohnologs, named as *crha* and *crhb*, have been identified (Grone & Maruska, 2015). The expression of *crha* has been mainly observed in the retina (Grone & Maruska, 2015; Kohei Hosono et al., 2015), with weak expression in the brain (about 100 times less than in retina) of fish (Kohei Hosono et al., 2015). On the other hand, the expression pattern of *crhb* was mainly characterized in the central nervous system (CNS), i.e., in the preoptic area, the hypothalamus, and the caudal neurosecretory system. For this reason, it has been related with the control of Acth in the pituitary gland (Alderman & Bernier, 2009; Bernier, Alderman, & Bristow, 2008; Carpenter, Maruska, Becker, & Fernald, 2014; Chen & Fernald, 2008; Grone & Maruska, 2015).

The action of CRH in the pituitary is mediated by the binding and activation of two highly conserved membrane receptors (CRH-R1 and -R2), which belong to class B of the G protein-coupled receptors (Lovejoy, Chang, Lovejoy, & del Castillo, 2014). Although in tetrapods, it has been reported that CRH has higher affinity for activate CRH-R1 (Vaughan et al., 1995), in teleosts, both Crhs have similar affinity for both Crh receptors (Kohei Hosono et al., 2015). Several studies in mammals have also demonstrated the ability of CRH receptors antagonists to block stress responses, such as anxiety or depression (Backstrom & Winberg, 2013; Grammatopoulos & Chrousos, 2002; Holsboer & Ising, 2008), placing CRH receptors at a critical point in regulation of HPA axis.

The molecular and morphological processes of masculinization by stress have been investigated at local, gonadal level, from nematodes (Christopher H. Chandler, Chadderdon, Phillips, Dworkin, & Janzen, 2012; C. H. Chandler, Phillips, & Janzen, 2008), fish (Hattori et al., 2007; Hayashi et al., 2010; Kitano, Hayashi, Shiraishi, & Kamei, 2012) and amphibians (M. Nakamura, 2009), to reptiles (Ge et al., 2018; Mork, Czerwinski, & Capel, 2014; Yatsu et al., 2015), but the involvement of the brain in sex-reversal is still under scrutiny. In all these vertebrates, exposure to environmental stressors during early life has several implications in reproduction. For instance, when reptiles and fish are exposed to stress during the critical period of gonadal differentiation, a strong bias in sex ratios can be induced (Capel, 2017; Fernandino, Hattori, Moreno Acosta, Strüssmann, & Somoza, 2013). The downstream factors involved in stress-induced masculinization in fish are well known (Hattori et al., 2009; Hayashi et al., 2010; Mankiewicz et al., 2013; Ribas et al., 2017; Tsalafouta et al., 2014; Yamaguchi, Yoshinaga, Yazawa, Gen, & Kitano, 2010), which in turn can act by three different mechanisms: (i) inhibition of estrogens synthesis (Kitano et al., 2012; Nozu & Nakamura, 2015), (ii) elevation of androgen synthesis (Fernandino, Hattori, Kishi, Strüssmann, & Somoza, 2012; Hattori et al., 2009), and (iii) apoptosis or meiotic arrest of germ cells (Yamaguchi & Kitano, 2012; Yamamoto et al., 2013). However, the molecular processes and key players controlling cortisol increase, that regulates these three mechanisms, remain unexplored.

In this study we provide clear evidence of the role of CNS in the regulation of the HPI axis, shedding light on the triggering mechanism of masculinization induced by environmental factors.

## RESULTS

### Expression of corticotropin-related genes reared at HT

First we examine the ontogeny of both *crh* paralogs regulation under normal and masculinizing temperature. The mRNA levels of *crha* and *crhb* were analyzed in medaka embryos at sages 26, 33 and 37, incubated at control (24 °C; CT) or high (32°C; HT) temperatures (Fig. 1). No differences were detected for *crha* between treatments, in any of the developmental stage (Fig. 1A). In contrast, we observed high transcript levels of *crhb* at HT for stage 37, corresponding to the gonadal sex determination period (Fig. 1B). Noteworthy, the expression levels of both *crha* and *crhb* were not affected by the sex genotype (XX vs XY) (Fig. S1).

**Fig. 1.**
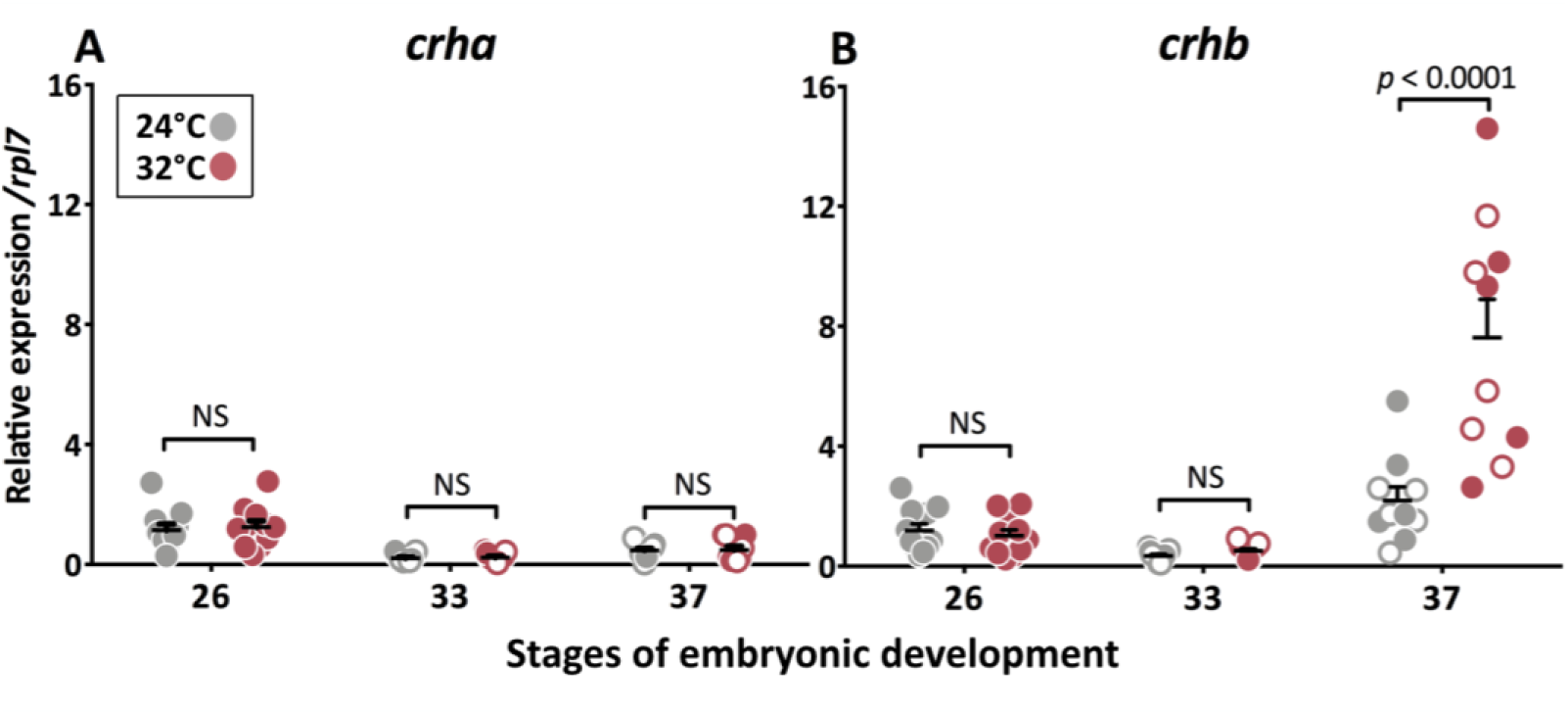
Developmental profiles of *crha* **(A)** and *crhb* **(B)** transcript abundance in embryos reared at 24 °C (control temperature) and 32 °C (high temperature). Data were measured by qPCR analysis in whole embryos at stages 26, 33, and 37. Gene expression levels are expressed relatively to the stage 26 group from 24 °C treatment. Quantification method was performed using the 2^-ΔΔCt^ method and values were normalized by the respective values of *rpl7*. Genotypic sex was determined at stages 33 and 37 by the presence/absence of the *dmy* gene; XX and XY are represented by filled circles and open circles, respectively. Horizontal bars indicate mean, with its respective standard error of the mean. The *p* values are indicated when transcript abundance between temperature treatments at the same developmental stage differ statistically (FgStatistics; *p* < 0.05).

Based in the up-regulation of *crhb* at stage 37 in embryos incubated at HT, we analyzed the transcript abundance of other HPI-related genes, such as *crh* receptors (*crhr1* and *crhr2*), the three urocortins, i.e., the urocortin1 (Ucn1)/sauvagine (Svg)/urotensin 1(Uts1), the urocortin2 (Ucn2), and the urocortin 3 (Ucn3) (Fig. 2A) (K. Hosono,Yamashita, Kikuchi, Hiraki-Kajiyama, & Okubo, 2017); and *acth* (Liu et al., 2003). Additionally, the expression of both *crhr*s, *crhr1* and *crhr2* (Fig. 2 E-F, respectively), was up-regulated at HT. No significant differences were observed in the transcript abundance of the other Crh-like genes, i.e., *uts1, ucn2*, and *ucn3* (Fig. 2B-D), and for *acth* (Fig. 2G), suggesting that these HPI axis-related genes are not regulated at the transcriptional level during exposure to thermal stress during early development.

**Fig. 2.**
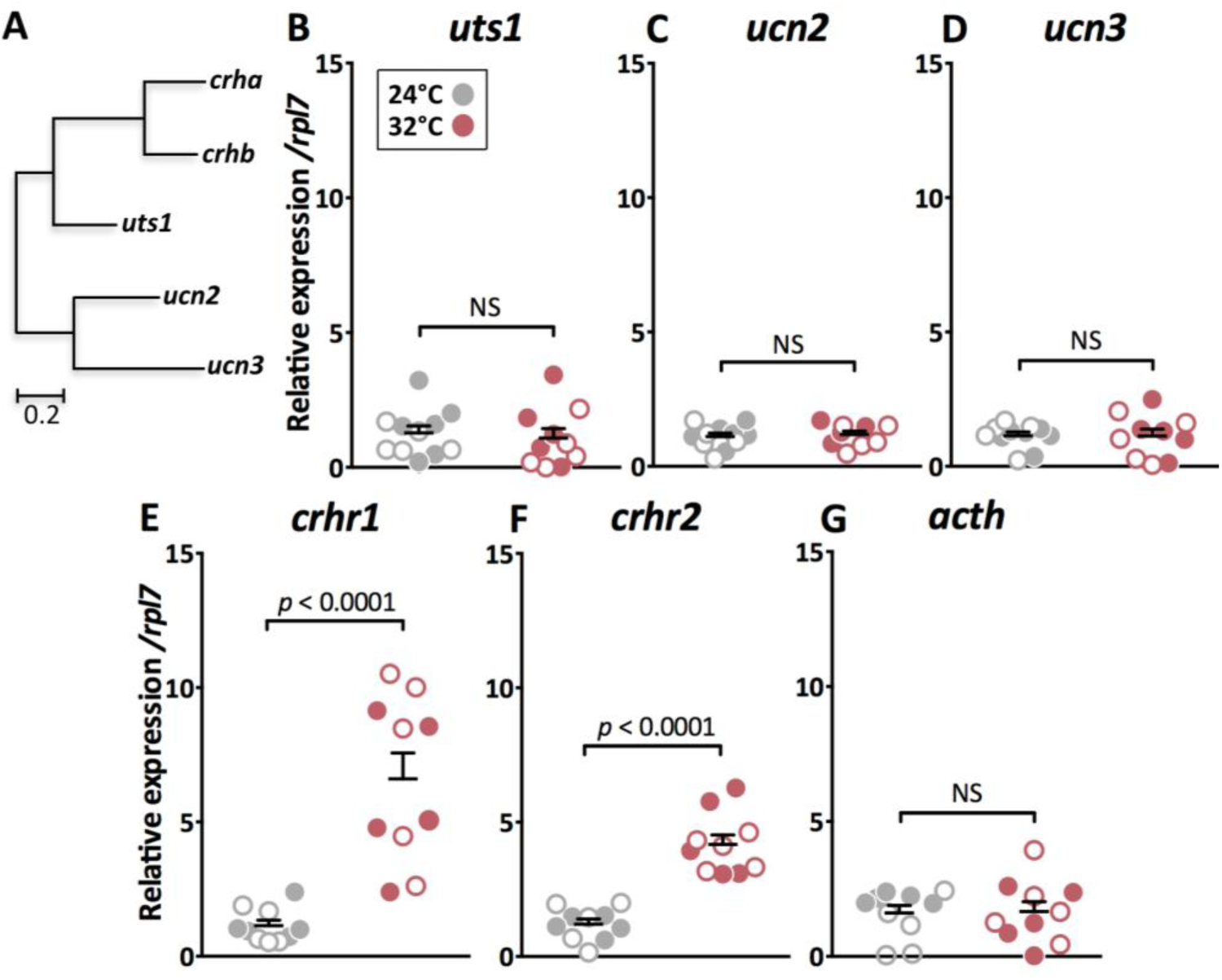
**(A)** Phylogenetic tree showing the relationship among Crh family peptides in medaka, obtained by Neighbor-joining method and a bootstrap test (MEGA 7.0 software). The scale beneath the tree reflects sequence distances. Genbank accession sequences are provided in Table 1. Gene expression profiles of urocortins **(B-D),** *crh* receptors **(E-F)**, *acth* **(G)**, and *gsdf* **(H)** in XX (filled circles) and XY (open circles) embryos reared at 24 °C and 32 °C. These data were measured by qPCR analysis in whole embryos at stage 37. Gene expression levels are expressed relatively to the 24 °C treatment. Quantification method was performed using the 2^-ΔΔCt^ method and values were normalized by the respective values of *rpl7*. Genotypic sex was determined by the presence/absence of the *dmy* gene, XX and XY are represented by filled circles and open circles, respectively. Horizontal bars indicate mean, with its respective standard error of the mean. The *p* values are indicated when transcript abundance between temperature treatments at the same developmental stage differ statistically (FgStatistics; *p* < 0.05).

### Generation of biallelic mutation of crhr1 and crhr2 using CRISPR/Cas9 technology

To analyze the participation of the HPI axis in temperature-induced masculinization, we disrupted this axis through the biallelic mutations of *crhr1* or/and *crhr2* using CRISPR/Cas9 technology. Biallelic mutations of both Crh receptors generated indels in the transmembrane domain resulting in a receptor with a protein segment that fails to anchor into the membrane lipid bilayer, and then unable to activate the intracellular G coupled protein (Grammatopoulos, 2012). Thus, the sgRNAs for *crhr1* and *crhr2* genes were designed at the exons 7 (located in the transmembrane helix 3) and 10 (located in the transmembrane helix 6; Fig. S2A), respectively. These sgRNAs were synthesized *in vitro*, and co-injected with *nCas9n* RNA (*cas9*) into one-cell-stage embryos. The mutagenesis efficiency for each sgRNA was analyzed by the heteroduplex mobility assay (HMA; Fig. S2B) (Ota et al., 2013), which reached 99.6 % for sgRNA-*crhr1* and 100 % for sgRNA-*crhr2* (Fig. S3A and S3B). Additionally, some biallelic positive amplifications were sequenced to confirm the indels presence (Fig. S2C). Data indicate that most cells contained biallelic indels, and consequently, loss of function in *crhr1* and *crhr2* mutants. Additionally, the potential off-target sites for each sgRNAs were searched in the medaka genome using the Medaka Pattern Match Tool (http://viewer.shigen.info/meda-kavw/crisprtool/) and CCTop - CRISPR/Cas9 target online predictor (Stemmer, Thumberger, del Sol Keyer, Wittbrodt, & Mateo, 2015). None of the embryos analyzed presented indels on the off-target sites for each of the injected sgRNAs (Fig. S3A and S3B).

Moreover, no morphological and survivals alterations were observed in a batch of animals reared at 24°C (CT) up to 60 days post-hatching (dph) in term of morphological and survival (Fig. S4).

### Genotypic female biallelic crhrs mutants did not show HT-induced masculinization

In order to assess the participation of Crh-related genes in the sex reversion of genotypic females to phenotypic males induced by HT, we analyzed the expression of well-known gene markers for gonadal sex differentiation in fish, such as *gsdf*, sry-box9 type 2α (*sox9a2*), gonadal aromatase (*cyp19a1a*, estrogen-related gene) and hydroxysteroid 11-beta dehydrogenase 2 (*hsd11b2*, androgen-related gene) (Chakraborty, Zhou,Chaudhari, Iguchi, & Nagahama, 2016; Fernandino et al., 2012; Imai, Saino, & Matsuda, 2015; Kurokawa et al., 2007; S. Nakamura et al., 2012; Shibata et al., 2010; X. Zhang et al., 2016; Zhou et al., 2016). Fertilized eggs were coinjected with *cas9* RNA and sgRNA for each of *crhr* (*cas9*+sgRNA-*crhr1* or *cas9*+sgRNA-*crhr2*) alone or together (*cas9*+sgRNA-*crhr1+*-*crhr2*); and they were then incubated at HT (32 °C). Control fertilized eggs were injected only with *cas9* and then incubated at CT and HT (*cas9*-24 °C and *cas9*-32 °C, respectively; Fig. 3A). In all treatments, genotypic females (XX, *dmy*^-/-^) that presented indels were selected for analysis of gene expression at stage 37. As expected, *cas9*-32 °C individuals presented higher levels of *gsdf* and *sox9a2*, and lower of *cyp19a1a* expression levels when was compared to *cas9*-24 °C individuals (Fig. 3B, 3C and 3D, respectively), evidencing the molecular mechanism of action of masculinization induced by HT. However, the double biallelic *crhr*s mutant of genotypic females at HT showed a female pattern of lower *gsdf* and *sox92a* expression levels, and higher of *cyp19a1a*, resembling those of *cas9*-24 °C group (Fig. 3B, 3C, and 3D).

When each biallelic *crhr* mutants of XX embryos were analyzed the gene expression pattern showed an intermediate phenotype, with high *gsdf, sox9a2* and *cyp19a1a* (Fig. 3B, 3C, and 3D). Here it is necessary to take into account that in the biallelic mutant of each *crh* receptor as the *crhr* paralog is fully active. Moreover, we analyzed the expression pattern of the androgen-related gene, *hsd11b2*, which did not show differences between treatments (Fig. 3F).

**Fig. 3.**
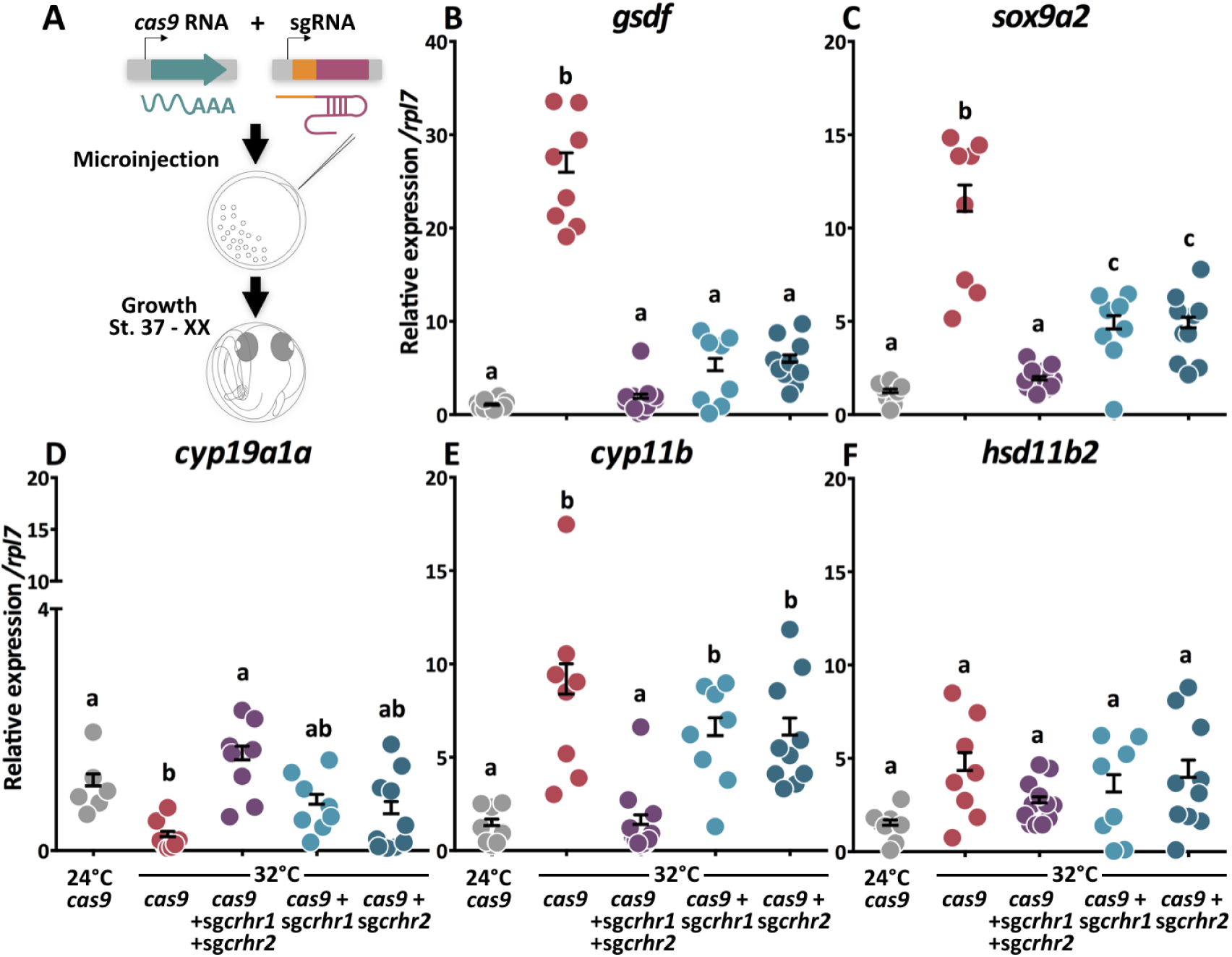
**(A)** Schematic representation of experimental procedure used to analyze the effect of loss function in *crh* receptors. **(B)** Transcript abundance profiles of testicular genes markers: *gsdf* and *sox9a2* **(B-C,respectively)**, the ovarian gene marker: *cyp19a1a* **(D),** interrenal gland gene marker *cyp11b* **(E),** and the androgen-related gene *hsd11b2* **(F)**, in the control (*cas9* injected embryos reared at 24 °C and 32 °C) and biallelic *crh* receptors mutants: *cas9*+sgRNA-*crhr1, cas9*+sgRNA-*crhr2* and *cas9+*sgRNA-*crhr1+*-*crhr2* coinjected embryos reared at 32 °C. These data were measured by qPCR analysis in whole embryos with genotypic sex XX at stage 37. Gene expression levels are expressed relative to *cas9* at the 24 °C treatment. Quantification method was performed using the 2^-ΔΔCt^ method and values were normalized by the respective values of *rpl7*. Horizontal bars indicate mean, with its respective standard error of the mean. Different letters indicate statistically significant differences between treatments (one-way ANOVA, followed by a Tukey’s multiple comparison test; *p* < 0.05).

### Crhrs are necessary to elicit sex reversion by high temperature

Besides the expression of testis and ovary-related gene markers, we also analyzed gonadal morphology of XX biallelic *crhr* mutants that were incubated at HT until hatching and thereafter at 26 °C (breeding temperature) for 20 dph, when gonad could be morphologically well differentiated. XX juveniles injected with *cas9* (control) and incubated at HT until hatching presented 68.8 % of sex reversion toward males, as evidenced by testis morphology (Fig. 4A and 4C). At 24 °C no reversal was found, with all fish showing normal ovary development (Fig. 4A and 4B). The double biallelic *crhrs* mutant showed a wide-ranging insensitivity to HT-induced female-to-male sex reversal, with all XX individuals presenting normal ovary morphology (Fig. 4A, and 4B). Moreover, in case of the biallelic *crhr1* mutant sex-reversed individuals were observed in 19 % of individuals (Fig. 4A and 4C). However, 19 % of intersex individuals were obtained, i.e., animals with ovaries containing spermatocytes (ova-testis; Fig. 4D). Finally, XX biallelic *crhr2* mutant juveniles showed 35% of sex reversal and 10 % of intersex gonads (Fig. 4A, 4C and 4D).

**Fig. 4.**
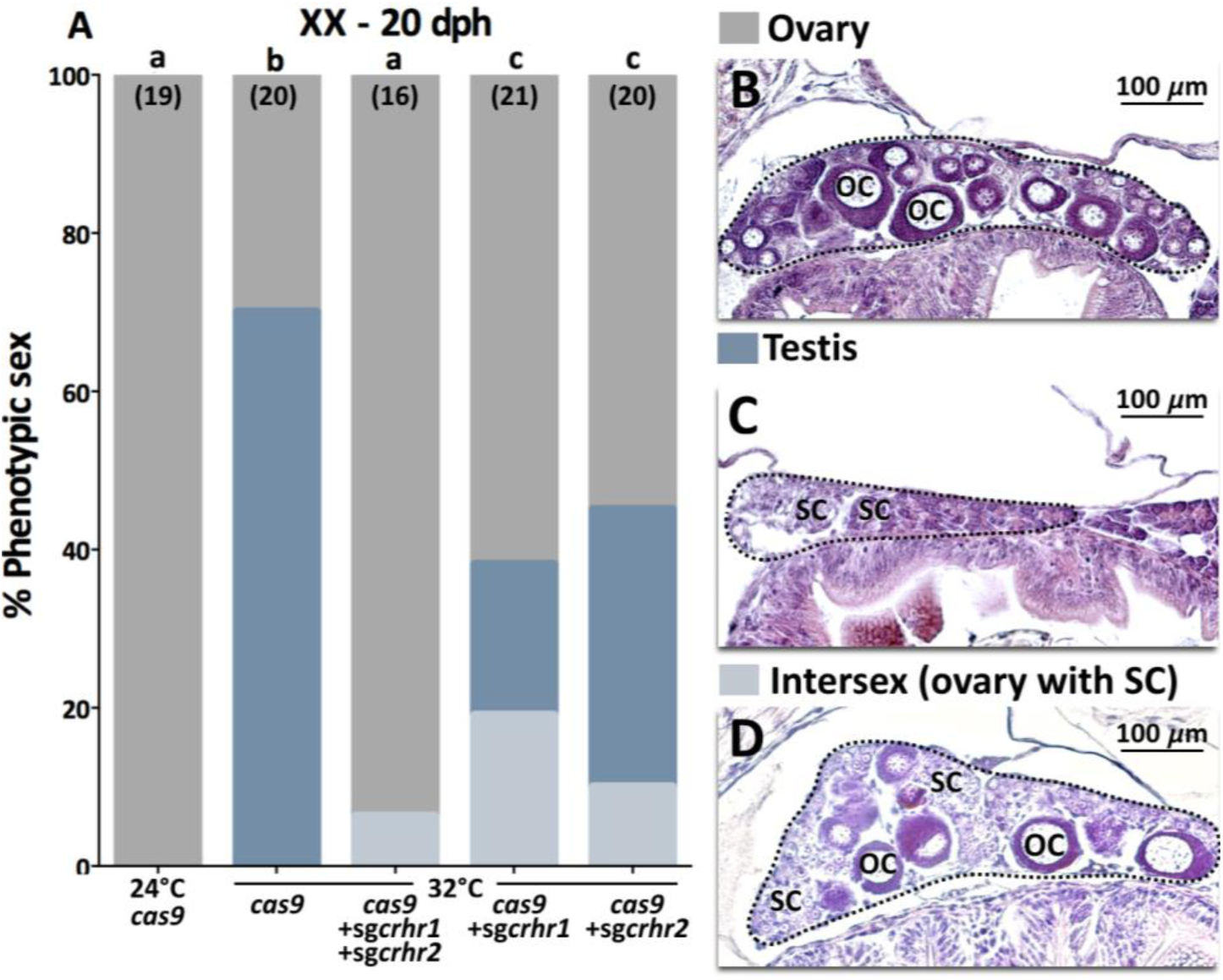
Participation of *crhr*s in the sex reversal induced by HT. **(A)** Percentages of genetic females (XX) with sex-reversed testicular morphology, **(B)** ovary (OC, oocytes), **(C)** testis with spermatocytes (SC), and **(D)** intersex (OC with SC) in embryos injected with *cas9* (control), *cas9*+sgRNA-*crhr1, cas9*+sgRNA-*crhr2* and *cas9+*sgRNA-*crhr1+*sgRNA-*crhr2.* The number of medaka juveniles analyzed in each treatment is shown between brackets. Different letters indicate statistically significant differences between treatments (one-way ANOVA, followed by a Tukey’s multiple comparison test; *p* < 0.05).

### Biallelic mutations of crhr exhibit inhibition of Acth release and lack of cortisol increase

As we previously did not observe a correlation between the up-regulation of *crhb* and the *acth* transcript abundance (Fig. 1, and 2), we measured the Acth-immunoreactive (Acth-ir) cells using immunofluorescence in the pituitary of genotypic female embryos at stage 39, with or without functional receptors and incubated them at HT (Fig. 5A). Firstly, we observed differences in the fluorescence intensity of the Acth-ir cells in XX embryos incubated at control and high temperature (Fig. 5B, 5C, and 5H), suggesting that thermal stress induces Acth release. Moreover, we measured Acth-ir in biallelic *crhrs* mutants and observed higher fluorescence intensity in relation to embryos incubated at HT (Fig. 5C, 5D, 5E, 5F, and S5), resembling the XX *cas9* control embryos (Fig. 5B, and S5). These results show that the biallelic mutation of *crhrs* in XX embryos causes the accumulation of Acth in pituitary cells, indicating that both *crh* receptors are mostly related to Acth release in stress response induced by high temperature.

To corroborate that biallelic mutations of *crhr*s do disrupt the HPI axis, the level of cortisol and the mRNA expression of P450 11-beta (*cyp11b*), enzyme expressed by the interrenal gland involved in cortisol synthesis (Montero et al., 2015), were measured in all treatment. We observed in both biallelic *crhr1* and *crhr2* mutants an increase of cortisol levels at the end of the gonadal sex determination period. On the other hand, the levels of cortisol in the double biallelic *crhr* mutant were completely suppressed (Fig.5H). Moreover, *cyp11b* was up-regulated at HT and down-regulated in the double biallelic *crhr*s mutant (Fig. 3E), showing that the gene involved in the synthesis of cortisol is transcriptionally active; *hsd11b2*, which is involved in cortisol catabolism and 11-oxygenated androgen synthesis, did not show differences between treatments (Fig. 3F).

**Fig. 5.**
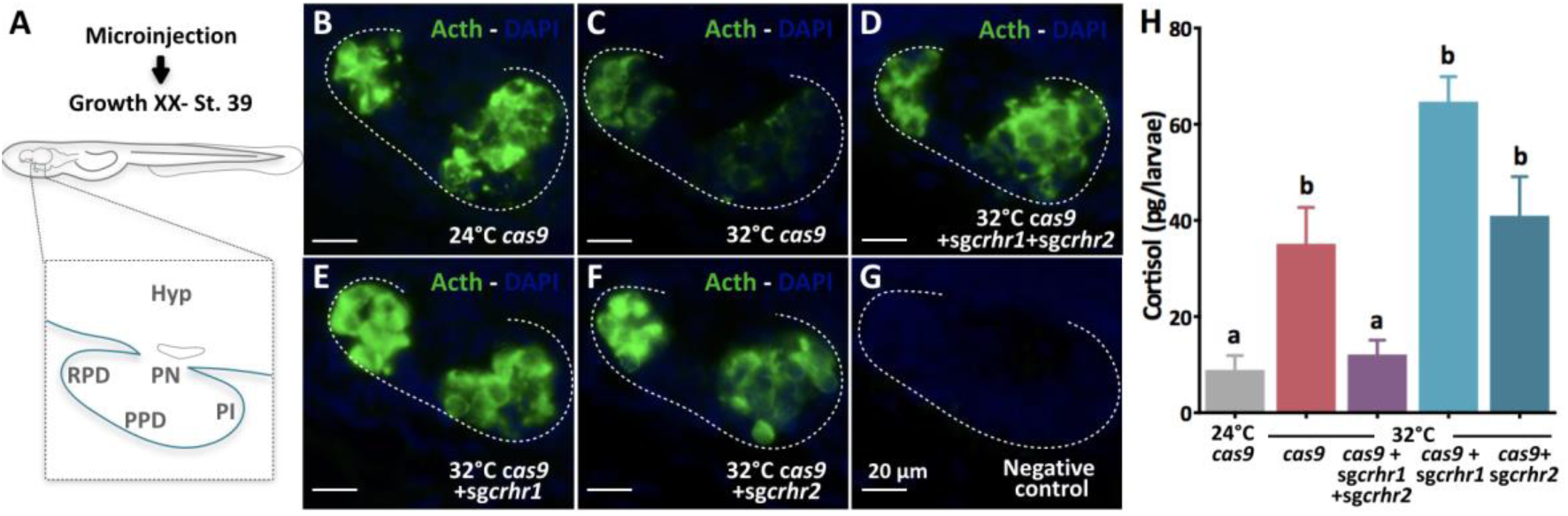
**(A)** Schematic representation of pituitary gland. Fluorescent images of Acth-ir cells in the pituitary of embryos injected with *cas9* RNA showing the negative control without **(B)** and with signal for Acth **(C)**, *cas9*+sgRNA-*crhr1* **(D)**, and *cas9*+sgRNA-*crhr2* **(E)** coinjected embryos reared at 32°C until stage 39. **(F)** Quantification of cortisol levels in embryos injected with *cas9* (controls at 24 °C and 32 °C), *cas9*+sgRNA-*crhr1, cas9*+sgRNA-*crhr2* and both *cas9+crh* receptors at 32 °C. Horizontal bars indicate mean, with its respective standard error of the mean. Different letters indicate statistically significant differences between treatments (one-way ANOVA, followed by a Tukey’s multiple comparison test; *p* < 0.05).

### Cortisol exposure rescued the lack of sex reversal phenotype in crhrs mutants

In view that the entire HPI axis seems to be functional during the critical period of gonadal fate and the biallelic mutations in *crhr*s inhibited masculinization of genotypic females incubated at HT, we decided to test whether the addition of cortisol could rescue the absence of sex reversal in the mutants. Therefore, we performed an experiment in which all embryos were maintained in an embryo medium with or without cortisol (5 μM) from fertilization to 5 dph (Fig. 6A) (Hayashi et al., 2010). The double biallelic *crhrs* mutants showed a transcription the phenotype of XX at HT, with a low transcript abundance of *gsdf* (Fig. 6B), a typical XX-24 °C *gsdf* expression pattern.

Finally, the treatments of XX biallelic cr*hr1* mutant, treated with or without cortisol at HT, presented high levels of *gsdf*, similar to control XX *cas9* injected larvae (Fig. 6B), suggesting that the XX biallelic *crhr1* mutant is not sufficient to induce a female (low) pattern of *gsdf*. These results are in agreement with the high level of cortisol observed at stage 39 (Fig. 5I). However, XX biallelic *crhr2* mutants larvae reared at HT maintained low transcript abundance of *gsdf* (Fig. 6B), a typical female-like expression pattern.

Most importantly, XX biallelic *crhr2* mutant reared with 5 μM cortisol at HT showed a male-like (high) *gsdf* expression pattern, similar to XX *cas9*-injected XX fish (Fig. 6B). To better understand the compensatory molecular mechanism, we analyzed the transcript abundance of the *crhr2* and *crhr1* in the XX biallelic *crhr1*and *crhr2* mutant, respectively. We also observed an up-regulation of the *crhr2* in the XX biallelic *crhr1* mutants (Fig. 6C), but not for *crhr1* in XX biallelic *crhr2* mutant larvae (Fig. 6D), suggesting a molecular compensatory mechanism.

**Fig. 6.**
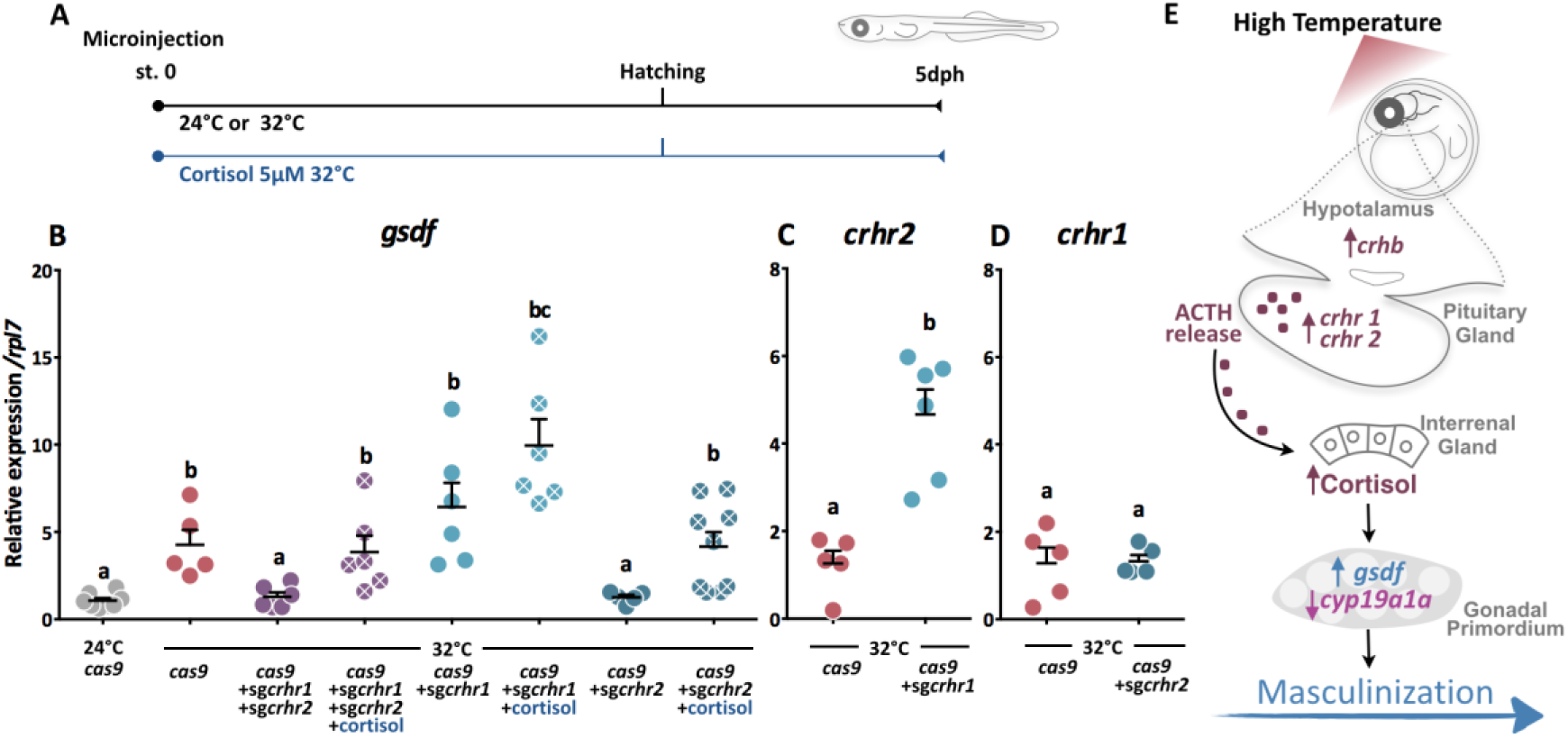
Rescue of masculinization in biallelic *crhr*s mutants phenotype by cortisol treatment. **(A)** Schematic representation of experimental procedure. **(B)** Gene expression profile of testicular gene marker *gsdf* was analyzed in control (*cas9*), *cas9*+sgRNA-*crhr1, cas9*+sgRNA-*crhr2*, and *cas9+*sgRNA-*crhr1+*sgRNA-*crhr2* treated with cortisol (5 μM) or vehicle (ethanol) between fertilization and 5 dph at 32 °C. Quantification of *crhr2* **(C)** and *crhr1* **(D)** expression in *cas9*+sgRNA-*crhr1* and -*crhr2* coinjected larvae, respectively. Gene expression levels are expressed relatively to *cas9*+sgRNA-*crhr1* without cortisol treatment (for panel **B**) and to *cas9* treatment (for panels **C** and **D**). Data were measured by qPCR in whole embryos with the genotypic sex XX. Quantification method was performed using the 2^-ΔΔCt^ method and values were normalized by the respective values of *rpl7*. Horizontal bars indicate mean, with its respective standard error of the mean. Different letters indicate statistically significant differences between treatments (FgStatistics; *p* < 0.05). **(E)** Schematic representation of the proposed mechanism by which the CNS, through the corticotropin releasing hormone b (*crhb*) and its receptors (*crhr1* and *crhr2*) are the transducer of stress-induced masculinization.

## DISCUSSION

Environmental factors that act during the critical period of fish gonadal development are able to alter sex ratios, especially toward males (Fernandino et al., 2013; Ospina-Álvarez & Piferrer, 2008). Even presenting established genotypic sex-determining mechanisms with known sex-determining genes, many fish species produce male-skewed sex ratios when environmental temperatures are elevated during early development. However, whether this phenomenon has any adaptive value or not is unknown for the vast majority of species. Although the understanding of this mechanism has great interest for basic biology and the perspective of global climate change, the pathways that mediate environmental cues and gonadal fate, and the involvement of extra-gonadal organs in this process are still unknown. Our results demonstrate for the first time the fundamental role of the CNS as the transducer in a form of environmental sex determination (ESD), through the regulation of the HPI axis.

In the current work we demonstrated that, during the gonadal sex determination period, the HPI axis is active. Moreover, we proved that out of two *crh* paralogs, only *crhb* was up-regulated at high masculinizing temperatures along embryonic development. In medaka, the *crha* gene was previously misidentified as a new member of Crh family and named as *telocortin* (*tcn*). The expression of *crha* has been mainly observed in the retina, with a weak expression in the brain (Kohei Hosono et al., 2015). In another teleost, *Astatotilapia burtoni*, the presence of *crha* has been related to the mediation of social information or stress responses in the visual system, facilitating signal processing before it even reaches the brain (Grone & Maruska, 2015). These previous results are in concordance with our observations, in which *crha* transcription does not seem to be induced by environmental stressors, such as high temperature. Besides *crha* and *crhb*, other members of the *crh* family genes are present in the medaka genome, such as the *uts1, ucn2,* and *ucn3* (K. Hosono et al., 2017). Known as urocortins, these genes code for neuropeptides that share structural similarity with *crh*s, and can act as additional endogenous ligands for CRH receptors. In mice, they have been involved in stress responses and also anxiety (Bale & Vale, 2004; Sztainberg & Chen, 2012). Nevertheless, neither of the urocortins was up-regulated during the sex determination period at high, masculinization temperature.

Regarding *crhb*, the high expression pattern at HT was in agreement with results of other well-known stress responses, at its role in regulating the release of glucocorticoids (Alderman & Bernier, 2009; Carpenter et al., 2014; Chen & Fernald, 2008; Grone & Maruska, 2015). Moreover, a similar pattern was obtained for others genes of the Crh-related pathway, crucial to the HPI axis is active (Lovejoy & de Lannoy, 2013), during the gonadal sex determination period, as for the *crh* receptors, *crhr1* and *crhr2*. In medaka, the first peak of cortisol occurs in 2 dph larvae, when animals are reared at normal breeding temperatures (Trayer, Hwang, Prunet, & Thermes, 2013). However, Hayashi et al. (Hayashi et al., 2010) and our study showed an early rise in cortisol in embryos reared at HT, at the time of the gonadal sex determination period, evidencing an earlier activation of mechanisms involved in the surge of cortisol levels. On this regard, the high expression of *crhb* and their receptors, *crhr1* and *crhr2*, in our study is consistent with the timing of cortisol increase. Notably, the overlapping between the timing of early activation of the HPI axis and the gonadal sex determination period is crucial to understand how high levels of cortisol are triggered and are related to male-skewed sex ratio (Hattori et al., 2009; Hayashi et al., 2010). Subsequently, in order to validate our hypothesis, we disrupted the HPI axis with biallelic mutation in both *crhr*s. These mutants were characterized by a lack of cortisol response at HT, down-regulation of testicular gene markers, and the concomitant inhibition of sex reversal (masculinization in XX) induced by stress. Thus, double biallelic *crhr*s mutants phenocopied the previous results on the inhibition of cortisol synthesis, with the absence of sex reversal in genotypic females (Hayashi et al., 2010). These observations corroborate for first time the participation of the brain in the stress-induced masculinization.

In all vertebrates, CRH regulates the synthesis and release of ACTH (Mommsen et al., 1999; Wendelaar Bonga, 1997) through their transmembrane receptors in the pituitary gland (Lovejoy et al., 2014). In the present work, although *acth* transcript abundance did not show any change during the gonadal sex determination period and under stress conditions, we detected low intensities of Acth-ir in HT embryos that could be explained by a stimulation of Acth release by thermal stress. Moreover, biallelic mutations of *crhr*s showed an accumulation of Acth in the pituitary, phenocoping the high fluorescence pattern of control group. Furthermore, these Acth accumulation or release are in concordance with the cortisol levels observed in each loss of *crhr*s function mutants. In mammals, both CRH and CRHR1 are associated with the HPA axis at the initial stress response whereas CRHR2 plays a major role during the chronic and later response to stress (Lovejoy & de Lannoy, 2013). CRHR1 knockout mice showed reduced stress-induced release of ACTH and corticosterone, providing evidence that CRHR1 mediates stress-induced hormone activation (Smith et al., 1998; Timpl et al., 1998). On the other hand, CRHR2-deficient mice possessed a generally normal initiation of the stress-response, but later on an early disruption of the ACTH release, suggesting that CRHR2 is also involved in the maintenance of HPA axis drive (Coste et al., 2000). In view of these considerations, our data are in accordance with results reported in mice, since the loss of function in each of receptors, *crhr1* or *crhr2*, or both together, resulted in a decrease of Acth release. Such disruption in HPI axis was demonstrated to be crucial for female-to-male sex reversal in our studies with medaka under high, stressful temperatures.

An in depth analysis on the molecular responses in each *crhr*s loss-of-function under thermal stress showed that, although embryos coinjected with *cas9*+sgRNA-*crhr1* or *crhr2* presented an early inhibition of *gsdf* expression, an increased cortisol level was observed at the end of the gonadal sex determination period when only one of the *crhr* was biallelically mutated. However, the level of cortisol in the double *crhr* biallelic mutant was completely suppressed. In each of biallelic *crhr* mutant is necessary taking into account that the paralog is fully active, explaining these partial compensation, and only half of sex reversal. In the second case, the strong decreased in cortisol level of the double *crhr*s mutants resembled the absence of stress response observed without an environmental stressor, with the concomitant absent of female sex reversal. In addition, the double *crhr* mutant phenocopied previous results observed by the cortisol synthesis inhibition in medaka (Hayashi et al., 2010), with the absence of sex reversal. Taken together, these results highlight the importance of the involvement of both Crh receptors in fish masculinization induced by environmental stressors.

Once the HPI axis has translated the stimulus of an environmental stressor, is important to know how cortisol transduces this response to masculinize the gonad. In some fish, including medaka, has been proposed that gonadal aromatase, an enzyme involved in estradiol synthesis, or other genes related to its regulation, such as FTZ-F1 - the ortholog of mammalian steroidogenic factor1 – are inhibited by cortisol (Hayashi et al., 2010; Navarro-Martin et al., 2011; Yamaguchi et al., 2010). Furthermore, in pejerrey (*Odontesthes bonariensis*) has been suggested that androgens, synthetized through the action of *hsd11b2*, are considered as mediators of stress (Fernandino et al., 2012; Fernandino et al., 2013). Our results confirm that *cyp19a1a* transcription is suppressed at HT, and demonstrated that high transcription levels can be rescued in double biallelic *crhr*s mutants.

Three different results, including (i) disruption of HPI axis, (ii) the increase of testicular gene markers with the concomitant decrease of sex reversal of genotypic females, and (iii) the rescue of masculinization with cortisol, support the fact CNS is involved in the sex reversal induced by environmental stressors (as summarized in the Fig. 6E), as contrasting to genotypic sex determination in which the sexual fate decision begins from the gonad.

## MATERIAL AND METHODS

### Source of animals and experimental conditions

Fertilized eggs of *O. latipes* were incubated in Petri disks of 70 mm with embryo medium (17 mM NaCl, 0.4 mM KCl, 0.27 mM CaCl_2_2H_2_O and 0.66 mM MgSO_4_; pH7) at 24 °C (CT) or 32 °C (HT). Sampling was performed at stages 26, 33, 37, 39, and at 5, 20 or 60 days after hatching (dah) (Iwamatsu, 2004). These stages corresponded to the end of primordial germ cells (PGCs) migration and the formation of the gonadal primordium (stage 26), the beginning of *dmy/dmrt1bY* transcription in gonadal somatic cells (stage 33), the sexual dimorphism in PGCs proliferation (stage 35-37), and to the maximum PGCs proliferation in XX embryos and latest embryo stage of the gonadal sex determination period (stage 39) (Saito et al., 2007). Based on previous work, we know that 5-dph larvae are sensitive to cortisol treatment (Hayashi et al., 2010), that 20-dph fish can easily be assessed for gonadal sex morphology, and that 60-dph animals have grown as adult fish to assess survivorship. The strain hi-medaka (ID: MT835) was supplied from the National BioResource Project (NBRP) Medaka (www.shigen.nig.ac.jp/medaka/). All fish were maintained and fed following standard protocols to medaka (M. Kinoshita, Murata, Naruse, & Tanaka, 2012). Fish were handled in accordance with the Universities Federation for Animal Welfare Handbook on the Care and Management of Laboratory Animals (www.ufaw.org.uk) and internal institutional regulations.

### RNA and quantification by RT-qPCR

Total RNA was extracted from individual embryos using the RNAqueos®-Micro kit (Ambion by Life Technologies) for stage 26, the Illustra RNAspin Mini was used for stage 33, and 350 μL of TRIzol® Reagent (Life Technologies) used for stage 37, 39, and 5 dph, following the manufacturer’s instructions. To perform the cDNA synthesis, RNA of each individual sample (250 ng) was treated with Deoxyribonuclease I Amplification Grade (Life Technologies) and reverse-transcribed using SuperScript II (Life Technologies) with oligo(dT) following the manufacturer’s instructions. Each primer pair was previously validated analyzing the melting curve, efficiency between 95-105%, with a slope of around -3.30 and a R2 value > 0.99. Real-time PCR primers are listed in Table S1. Samples were analyzed with Step One Plus Real-Time PCR System (Applied Biosystems). The amplification protocol consisted of an initial cycle of 1 min at 95 °C, followed by 10 s at 95 °C and 30 s at 60 °C for a total of 45 cycles. The subsequent quantification method was performed using the 2^-ΔΔCt^ method (threshold cycle; www.appliedbiosystems.com/support/apptech) and normalized against reference gene values for ribosomal protein L7 (*rpl7*) (Z. Zhang & Hu, 2007).

### Sexing of embryos by PCR

Each embryo of stages 33, 37, 39, and 5-dph and 20-dph larvae was analyzed to determine its genotypic sex. Animals were subjected to DNA analysis for the presence of the *dmy/dmrt1bY* gene. For this purpose, we collected DNA from each RNA extraction following manufacturer’s instructions. A PCR analysis was then performed using primers for *dmy* (Nanda et al., 2002) and the presence of *β-actin* gene was used as a DNA loading control (Table S1) (Hattori et al., 2007). The PCR products were analyzed on a 1% agarose gel.

### CRISPR/Cas9 target site design and single guide RNA (sgRNA) construction

CRISPR/Cas9 target sites were designed using the CCTop - CRISPR/Cas9 target online predictor (crispr.cos.uni-heidelberg.de/index.html)(Stemmer et al., 2015), which identified sequence 5′ GG-(N18)-NGG3′ in exon 7 of *crhr1* (TTGAGGAACATCATCCAC TGG) and exon 10 of *crhr2* (GAGGCAGCAAGACGAGTG TGG) (Fig. S2A and S2C). Each sgRNA was synthesized by cloning the annealed oligonucleotides into the sgRNA expression vector pDR274 (Addgene #42250) (Hwang et al., 2013) followed by *in vitro* transcription, previously established by Ansai and Kinoshita (Ansai & Kinoshita, 2014). Briefly, a pair of oligonucleotides at final concentration of 10 mM each was annealed in 10 mL of annealing buffer (40 mM Tris-HCl [pH 8.0], 20 mM MgCl_2_, and 50 mM NaCl) by heating to 95°C for 2 min and then cooling slowly to 25°C. Then, the pDR274 vector was digested with BsaI-HF (New England Biolabs), and the annealed oligonucleotides were ligated into the pDR274 vector. The sgRNA expression vectors were digested by DraI, and the sgRNAs were synthesized using the MEGAshortscript T7 Transcription Kit (Thermo Fisher Scientific). The synthesized sgRNAs were purified by RNeasy Mini kit purification (QIAGEN). These RNA sequences were diluted to 50 ng/μL.

### Capped Cas9 RNA synthesis

The capped *cas9* (nCas9n RNA) was transcribed from pCS2-nCas9n plasmid (Addgene #47929). First, the plasmid was linearized by NotI and capped *cas9* was synthesized by mMESSAGE mMACHINE SP6 kit (Life Technologies). The synthesized *cas9* was purified by RNeasy Mini kit purification (QIAGEN). These RNA sequences were diluted to 200 ng/μL.

### Microinjection into embryos

Microinjection was performed into fertilized medaka eggs before the first cleavage as described previously (Masato Kinoshita, Kani, Ozato, & Wakamatsu, 2000). For CRISPR/Cas9 system, 25 ng/μl *sgRNA* and 100 ng/μL *cas9* were coinjected in 4.6 nL of RNA mixture. Embryos injected with *cas9* were used as controls. Microinjection was performed with a Nanoject II Auto-Nanoliter Injector (Drummond Scientific) coupled to a stereomicroscope (Olympus).

### DNA extraction to Heteroduplex mobility assay (HMA)

To analyze the efficiency and specificity of the CRISPR/Cas9 system, 3 days post-fertilization embryos were used (Ansai & Kinoshita, 2014). Genomic DNA was extracted by incubating each medaka embryo in 25 μL of 5 mM NaOH, 0.2 mM EDTA at 95°C for 5 min. After cooling to room temperature (RT) 25 μL of 40 mM Tris-HCl, pH 8.0, was added to the extract. The supernatant was used as template for PCR to HMA. Conventional PCR analysis was performed with genomic DNA using primers listed in Table S1. Electrophoresis performed in 12 % acrylamide gel (Ota et al., 2013), stained with ethidium bromide for 15 min before examination. PCR products were sequenced to confirm the presence of indels (Ansai & Kinoshita, 2014).

### Off-target analysis

Potential off-target sites in the medaka genome were searched using a “Pattern Match” tool in New Medaka Map (beta) at the NBRP medaka web site (http://viewer.shigen.info/medakavw/patternmatch). All potential off-target sites identified were analyzed by HMA using the primers listed in Table S1.

### Biallelic mutant screening

Crispant (injected embryos with *cas9*+sgRNA) fish were mated with wild-type ones from Himedaka strain. Genomic DNA was extracted from each F1 embryos for analysis of mutations by HMA, as described previously (Table S1). Mutant alleles in each embryo were determined by direct sequencing of the *crhr1* or *crhr2* gene region.

### Histological analysis

Samples for histological examination of gonadal sex (n = 15 – 25/per group) were taken at 20 dph and analyzed following the criteria reported above (5). Firstly, the caudal fin was taken for gDNA extraction using conventional saline buffer extraction to determine genotypic sex and for HMA analysis (Aljanabi & Martinez, 1997). The body trunk was fixed in Bouin’s solution and processed according to standard protocols for preparation of hematoxylin-eosin stained histological sections. These preparations were examined under the Nikon ECLIPSE Ni-U microscope (Nikon) and captured with a Digit Sight DS-Fi2 digital camera (Nikon).

### Immunofluorescence analysis of Acth

Medaka embryos at stage 39 from the different treatments were used. All individuals were processed under the same condition for fixation, washing and incubation with serum and antibody. The stage 39 was chosen to analyze the release of Acth upon up-regulation of *crhb*, which was detected at stage 37. The tail was used for sex genotying by PCR and HMA analysis and the rest of the body was fixed in Bouin’s solution overnight. Sections were then washed with 0.1 M phosphate-buffered saline (PBS pH 7.4) and blocked in 0.1 M PBS containing 0.5% of bovine serum albumin (Sigma-Aldrich) for 60 min before overnight incubation with a mixture of primary antibody against ACTH-NIDDK-anti-hATCH-IC-3 (rabbit, 1:250; kindly provided by Dante Paz, Universidad de Buenos Aires) at RT. After incubation, the sections were washed twice in PBS for 10 min each and incubated at RT for 90 min with the secondary antibody goat-anti-rabbit IgG (Life Technologies) conjugated with Alexa Fluor 488 (green), at a dilution of 1:2000 in PBS. Separate sets of slides were treated only with secondary antibody (negative controls). After incubation, sections were rinsed twice with PBS and mounted with mounting medium Fluoromount (Sigma Aldrich) containing 4’,6-diamidino-2-phenylindole (DAPI, 5 μg/ml, Life Technologies). Section photographs were taken using the Nikon Eclipse E7000 and the Image Pro Plus (Media Cybernetcs) at same capture conditions of exposure and gain to all samples. Finally, images were analyzed and measured for fluorescence using ImageJ (https://imagej.nih.gov/ij/) employing the relation of the fluorescence intensity of the area of pituitary gland and the mean fluorescence of background.

### Levels of cortisol

Enzyme immunoassay (EIA) was performed using the Cortisol Express EIA Kit according to instructions from the manufacturer (Cayman Chemical, Ann Arbor, USA) and previously used by our group (Fernandino et al., 2012). Briefly, pools of 23-25 embryos both sexes were immediately frozen at -80 °C, homogenized in 0.2 mL of PBS, and used for steroid extraction with 1 mL of diethyl ether. This procedure was repeated two times. After evaporation of the diethyl ether, samples were immediately resuspended in 2 mL EIA buffer and analyzed in a microplate reader (Rayto Model RT-2100C, Hong Kong) following the kit instructions. The recovery rate was estimated by the cold-spike method to be 0.85% and the intra- and inter-assay variation (CV%) ranged from 4 to 13%.

### Rescue of biallelic mutant phenotype by cortisol treatment

Both *cas9+*sgRNA-*crhr1* and/or +sgRNA-*crhr2* coinjected fish were treated with 5 μM of cortisol (18) (11β-11,17,21-trihydroxypregn-4-ene-3,20-dione; Sigma-Aldrich) from fertilization to 5 dph. Briefly, after the injection with a mixture of sgRNA (*crhr1* and/or *crhr2*) and *cas9*, the embryos were placed in Petri dish of 70 mm with embryo rearing medium (25 mL) supplemented with cortisol or vehicle control (with the same volume of stock solvent: 4.53 μL ethanol, 0.018%). The medium was changed every day. Both groups were reared at HT.

### Statistical analysis

All values are presented as mean ± standard error of the mean (SEM). Fold change and statistical analysis of RT-qPCR quantifications were performed by using FgStatistics interface (http://sites.google.com/site/fgStatistics/), based on the REST method from Pfaffl et al. (Pfaffl, Horgan, & Dempfle, 2002). The immunohistochemistry quantification was analyzed using χ^2^-distribution and statistical analyzes were performed using SPSS v20 program, using one-way Analysis of Variance (ANOVA), followed by a Tukey’s multiple comparison test. The differences on sex ratio were analyzed with the Hypothesis Testing to Compare Two Population Proportions. All statistical differences were accepted as significant when *p* < 0.05.

## ACKNOWLEDGMENTS

We thank Gabriela C. López (INTECH) for helping with histological and immunohistochemical preparations. We also thank Masato Kinoshita (Kyoto University) for teaching and helping with CRISPR/Cas9 technique, Adrián Mutto (Insituto de Investigaciones Biotecnológicas-UNSAM) for helping with microinjections, and Ricardo S. Hattori (Unidade de Pesquisa e Desenvolvimento de Campos do Jordão, APTA/SAA) for helpful advice. We are grateful to NBRP Medaka (https://shigen.nig.ac.jp/medaka/) for providing hi-medaka (Strain ID: MT835).

## THIS WORK WAS SUPPORTED BY GRANTS FROM

Consejo Nacional de Investigaciones Científicas y Técnicas, Fellowship Program for partial financing for a short training period D2979/16 (to J.I.F.), Agencia Nacional de Promoción Científica y Tecnológica Grants 1565/14 and 2501/15 (to J.I.F.), and 2783/15 (to G.M.S.). This work was also supported by a Discovery Grant from the Natural Sciences and Engineering Research Council (NSERC) of Canada (RGPIN 418576-2012) and a Canada Research Chair (CRC) to VSL. DCCC and LFAP were supported by a PhD scholarship from the National Research Council (CONICET). JIF and GMS are members of the career of scientific researcher at the CONICET. DCCC and LFAP are Ph.D. scholarship from CONICET.

## ADDRESS ALL CORRESPONDENCE AND REQUESTS FOR REPRINTS TO

Juan Ignacio Fernandino, Av. Intendente Marino Km. 8.2 (B7130IWA). Chascomús, Provincia de Buenos Aires, Argentina.

